# Identification of metabolites from tandem mass spectra with a machine learning approach utilizing structural features

**DOI:** 10.1101/573790

**Authors:** Yuanyue Li, Michael Kuhn, Anne-Claude Gavin, Peer Bork

## Abstract

Untargeted mass spectrometry is a powerful method for detecting metabolites in biological samples. However, fast and accurate identification of the metabolites’ structures from MS/MS spectra is still a great challenge. We present a new analysis method, called SF-Matching, that is based on the hypothesis that molecules with similar structural features will exhibit similar fragmentation patterns. We combine information on fragmentation patterns of molecules with shared substructures and then use random forest models to predict whether a given structure can yield a certain fragmentation pattern. These models can then be used to score candidate molecules for a given mass spectrum. For rapid identification, we pre-compute such scores for common biological molecular structure databases. Using benchmarking datasets, we find that our method has similar performance to CSI:FingerID and that very high accuracies can be achieved by combining our method with CSI:FingerID. Rarefaction analysis of the training dataset shows that the performance of our method will increase as more experimental data become available.

## Introduction

Untargeted mass spectrometry is a common approach for identification of metabolites in biological samples (Beger et al., 2016; O’Kell et al., 2017; Schrimpe-Rutledge et al., 2016). Thereby, a complex biological sample is analyzed with liquid chromatography electrospray ionization tandem mass spectrometry (LC-ESI-MS/MS), generating several thousands of MS/MS spectra in a few minutes. Inferring all molecular structures from these spectra in a fast, precise manner is, however, still a challenge. The currently fastest way of analyzing such data is to match fragmentation spectra of unknown substances to a reference spectral library (Kind et al., 2018). These spectral libraries are usually built from known purified metabolites, analyzed by mass spectrometry. Some databases like XCMS (Benton et al., 2015), GNPS (Wang et al., 2016) and Massbank (Horai et al., 2010) are collecting these data. This experimental approach has the highest accuracy, however, generating these reference libraries is money- and time-consuming.

*In silico* fragmentation methods, such as MS-FINDER (Tsugawa et al., 2016), MetFrag (Ruttkies et al., 2016) and CFM-ID (Allen et al., 2014) strive to explain all fragment ions; these methods break every possible covalent bond, scoring each broken bond based on its strength. Some methods like CFM-ID can generate a pre-calculated spectral library. However, as molecular rearrangements occur during fragmentation, precise prediction of the rearrangement is very difficult, and these methods suffer from a low identification accuracy (Blaženović et al., 2018). Other approaches like CSI:FingerID (Dührkop et al., 2015; Ludwig et al., 2018) convert a spectrum into a fragmentation tree, search this fragmentation tree against a database of known trees, and then infer a molecular fingerprint. This method shows high accuracy; however, it needs to search fragmentation trees one by one in an online way, which makes it time-consuming when analyzing many spectra.

Depending on the type of biological sample, different chemical search spaces can be used. For well-studied sample types, such as cultured cells or human plasma, many of the metabolites in these samples have been analyzed before. In these cases, spectra can often be matched to reference libraries. Even if reference spectra are not available, there is a high chance that the compounds are part of curated databases such as the KEGG pathway database (Kanehisa et al., 2017) or the HMDB database of metabolites (Wishart et al., 2018). For such compounds, several identification methods have be developed (Blaženović et al., 2018). More complex samples such as those from plants or the environment contain many molecules whose structures have not been determined yet, hampering compound identification using MS.

Here, we describe a new method called SF-Matching (<u>S</u>ub<u>F</u>ragment-<u>Matching</u>) to predict likely peaks in tandem mass spectra for small molecules using a machine learning approach. Circumventing the complexities of accurately modeling the fragmentation processes and probabilities of bond breakage, our new approach relies on detecting “fragile” substructures in the molecule. These enable us to derive the respective fragmentation patterns to achieve high identification accuracy of compounds from mass spectra. Furthermore, our approach appears complementary to the existing method CSI:FingerID as a combination with it achieves a much higher accuracy than either method on its own with only a small sensitivity decrease.

## Results

Our concept assumes that molecules consist of fragile and relatively stable substructures. Molecules with similar fragile structures will share similar fragmentation patterns even if they have different stable substructures. For example, in lysophosphatidylinositol and phosphatidylethanolamine, the inositol moiety, ethanolamine and the alkyl chain are stable substructures, connected by a fragile substructure containing ester bonds that are likely to lead to fragmentation. During fragmentation, the two different molecules with different alkyl chains will generate similar fatty acids as fragment ions, although the masses of their fragment ions are different (Figure 1a). In the fatty acid that can be detected as a fragmentation product, the alkyl chain is the stable substructure and the carboxyl group is the fragment of the fragile substructure.

**Figure 1:**
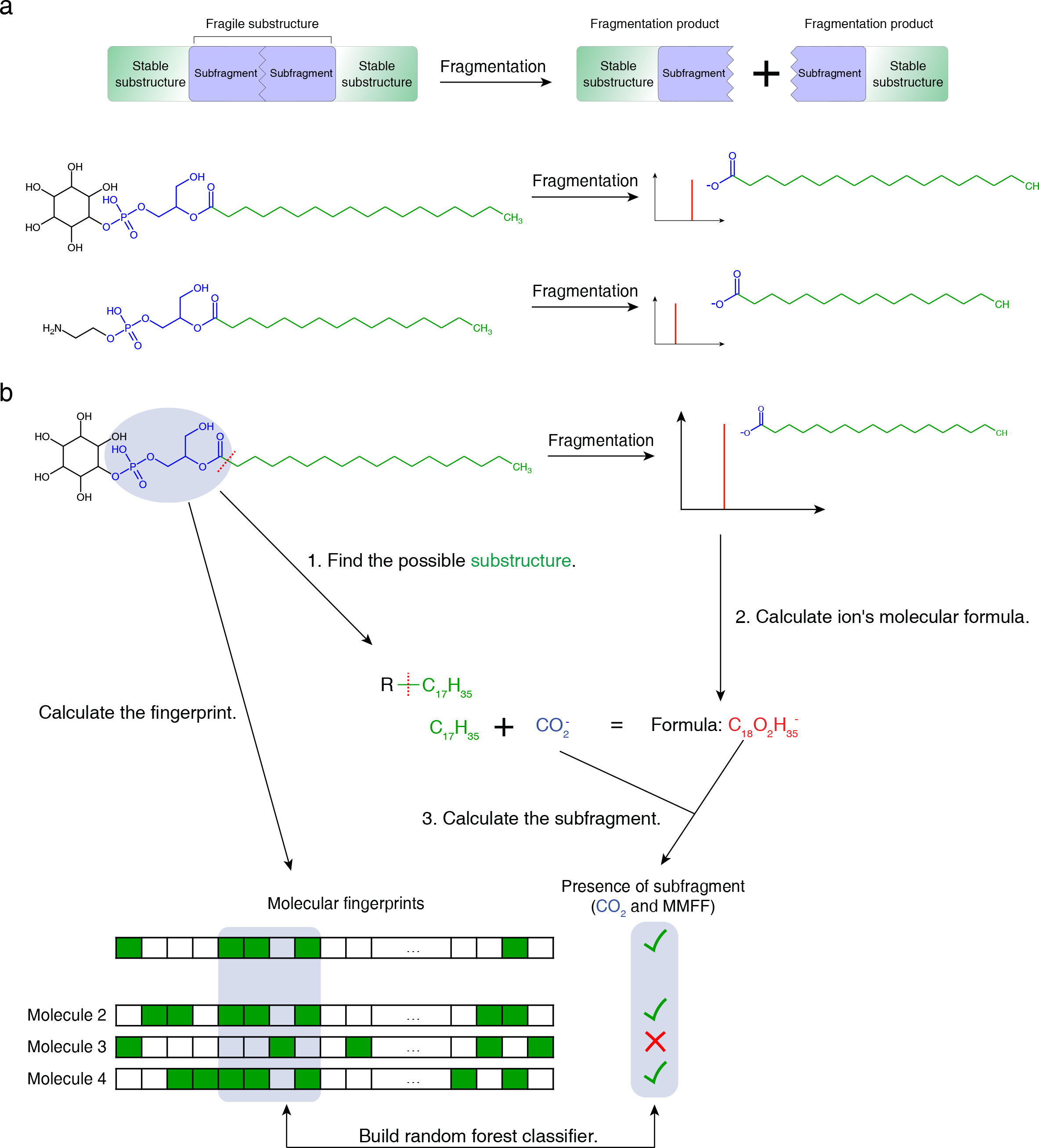
Schematic of the method. (a) The fragmentation of a molecule can be considered as breaking a fragile substructure surrounded by stable substructures. The resulting fragment contains both a stable substructure and part of the fragile substructure (which we term “subfragment”). (b) For training the method, we compare possible substructures to peaks in a reference database and calculate the subfragments that were observed. The presence of such a fragile subfragment can then be predicted based on the molecular fingerprint of the complete molecule.

Our aim was therefore to develop a method that can detect the presence of fragile substructures and based on this predict if a given spectrum is likely to belong to a particular compound. Unlike previous approaches, we use machine learning to associate structural information contained in 2D chemical fingerprints with fragmentation patterns. Molecular fingerprints encode the presence of various substructures of the molecule in a vector of bits. If two molecules have similar fragile substructures, some bits of the fingerprint will be the same. As the substructure may vary from molecule to molecule, in our approach, we do not try to identify the extract fragile substructure, but instead use the fingerprint to represent it, and use machine learning to detect the predictive parts of the fingerprint. Given a training database of molecular structures and mass spectra, we process each spectrum individually. On a high level, training the model works as follows (Figure 1b):

1. In the molecule associated with the spectrum, we find all substructures by individually breaking all covalent bonds in the molecules. For each broken bond, we record the resulting molecular formulas together with the bond type of the broken bond. For example, -C_17_H_35_ is one of the substructures of lysophosphatidylinositol.
2. For all peaks in the spectrum, we calculate the fragmentation products’ molecular formulas based on their m/z.
3. For each fragmentation product, we check which substructures found in step 1 are a subset of the fragmentation product’s molecular formula. For any fragmentation product, there may be several such substructures. Considering each substructure individually, we then designate it as the stable substructure. The remaining part of the fragmentation product must therefore be a part of the fragile substructure, and its molecular formula can be determined from the formulas of the fragmentation product and the stable substructure. Together with the bond type information, we call this part of the fragile substructure a “subfragment.” Each molecule in the database can be checked if it contains a subfragment.

For each known molecule, we calculate its molecular fingerprint based on the structure of the complete molecule. After determining the presence of subfragments across all training spectra, random forest classifiers are trained for each subfragment based on the molecular fingerprint and the presence of the subfragment in each spectrum.

For testing if a certain molecule is likely to belong to a given spectrum, we do the reverse: we calculate its molecular fingerprint and find all possible subfragments based on the peaks of the spectrum. Then for every subfragment, we use the corresponding random forest classifier to predict the probability of the subfragment. After finding the possible molecular substructures, we can calculate the mass of all possible fragment ions by adding the formula of subfragment to the formula of substructure.

To increase the speed of the method, we pre-calculated the predicted spectra of all biomolecules in four databases that collect most of the known, relevant biological molecules (KEGG (Kanehisa et al., 2017), HMDB (Wishart et al., 2018), ChEBI (Hastings et al., 2016) and ChEMBL (Gaulton et al., 2017)). This allows identification of compounds at a rate of more than 10 spectra per second in laptop with SSD hard disk. The pre-calculated database and searching scripts can be downloaded from http://www.bork.embl.de/Docu/sf_matching.

We evaluated the performance of our approach by comparing it with CFM-ID (Allen et al., 2014), and CSI:FingerID (Dührkop et al., 2015), the two methods that showed very good performance in CASMI 2016 and can also run in batch mode (Schymanski et al., 2017). To achieve a fair comparison, we removed all spectra from the test set that were part of our training dataset. First, we used all spectra provided in the context of the CASMI 2016 and 2017 automated structural identification challenge (Schymanski et al., 2017) as benchmark dataset. To estimate the performance on multiple chemical databases, we limited the candidates to the molecules that are in the selected database. If one molecule was not present in the target database, the corresponding spectrum was not considered. In the CASMI 2016 dataset, SF-Matching had the best performance when searching against four different databases of known molecules (Figure 2a). In the CASMI 2017 dataset, the performances of all methods dropped to less than 60%. SF-Matching performed slightly worse than CFM-ID for three of the chemical databases (Figure 2b). For ChEMBL, SF-Matching had much lower performance. As an additional benchmark, we also evaluated the methods using spectra from the EMBL Metabolomics Core Facility (EMBL-MCF) spectral library (Palmer et al., 2018). We selected candidate molecules from the respective chemical database with an m/z within 5 ppm of target molecules. In this dataset, SF-Matching had a similar performance as CSI:FingerID and was superior to CFM-ID (Figure 2c).

**Figure 2:**
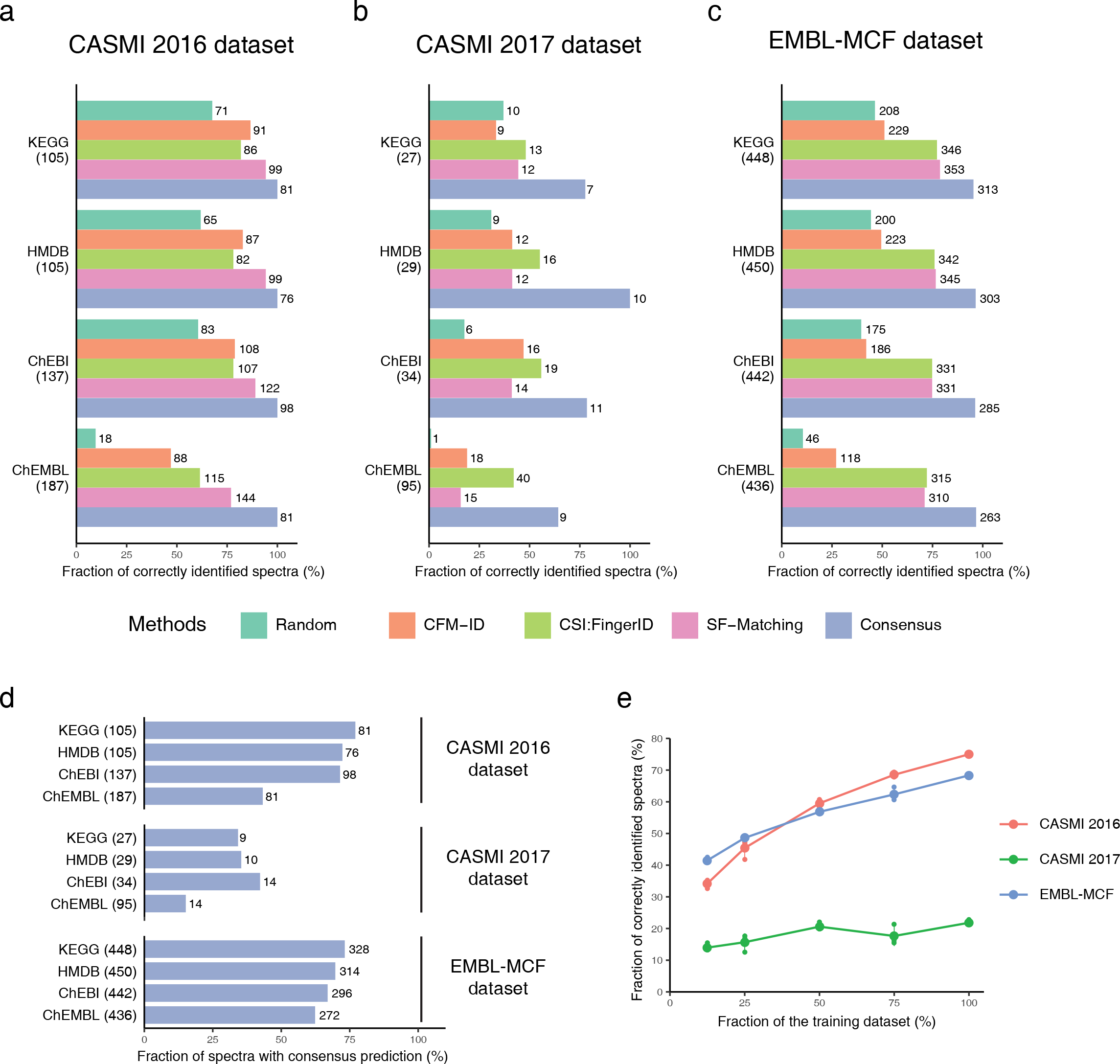
Performance evaluations. The accuracy and the sensitivity of SF-Matching, and the consensus method is compared against random prediction and two established methods on (a) the CASMI 2016 dataset, (b) the CASMI 2017 dataset, and (c) the EMBL-MCF dataset. The number in the parentheses indicates the total number of molecules which are contained in the various chemical databases; on right of the bar the number of correctly identified molecules is shown. (d) shows the fraction of spectra with consensus prediction. The number in the parentheses indicates the total number of molecules which are contained in the various chemical databases; on right of the bar the number of spectra with consensus prediction is shown. (e) shows the effect of training dataset size on the performance on all datasets. For each training dataset size, three training datasets were randomly sampled from the origin whole dataset, here we use the combination of four database as candidate database.

As our concept differs from existing ones, we reasoned that it should be possible to achieve a better accuracy if we combine prediction methods. As CSI:FingerID showed good performance in all the three benchmark dataset, we selected spectra where both our method and CSI:FingerID gave the same results. These consensus results achieved about 20% increase in accuracy than any single method, reaching more than 90% accuracy when analyzing the CASMI 2016 and EMBL-MCF datasets, and still more than 70% when analyzing the CASMI 2017 dataset. Due to the consensus calculation this comes at the cost of making predictions for fewer spectra, in average, around 55% spectra had a consensus identification. The fraction varied between 40%-80% in the CASMI 2016 and EMBL-MCF datasets and between 10%-45% in CASMI 2017 (Figure 2d).

As machine learning approaches gain power with increasing training sets, we randomly selected subsets of the training dataset to evaluate the performance. Indeed, we observed an increase in the prediction performance with increased training set size (Figure 2e). This result suggests that SF-matching will increase performance with time with more experimental spectra becoming available.

Taken together, we have developed a new method called SF-Matching to identify the spectra of small molecules in biological samples. Depending on the goals of the MS experiments, SF-matching itself can be used to contribute to candidate molecule identification, given its stand-alone performance, but it can also be used in combination with CSI:FingerID for candidate predictions with high accuracy. Furthermore, as expected from machine learning techniques, the power of the method will increase in the future with the addition of diverse known spectra of biomolecules.

## STAR Methods

### Converting spectra to subfragments

Given a molecular structure, we search for molecular substructures first. We remove all possible bonds *b*_*s,t*_ connecting atoms *a*_*s*_ and *a*_*t*_, with the exception of carbon–carbon bonds in the aliphatic chain and bonds within ring systems. Removing a bond yields two sub-molecules with molecular formulas *F*_*s,t*_ and *F*_*t,s*_. For each heavy atom *a*_*i*_ in a molecule, the Merck Molecular Force Field (MMFF) (Halgren, 1996) atom type 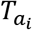 was determined and the bond type 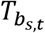 was defined as 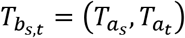.

The molecule’s corresponding spectrum can be treated as a list of n peaks {*P*_1_, *P*_2_,…, *P*_*n*_}, here *P*_*i*_ = (m/z *M*_*i*_, intensity *I*_*i*_), and the formula of a fragment ion *F*_*i*_ can be determined from *M*_*i*_. Then, we test for all *F*_*s,t*_ if they are a subset of *F*_*i*_. In this case, we can calculate the difference of the molecular formulas Δ_*i,s,t*_= *F*_*i*_ − *F*_*s,t*_, and define the subfragment 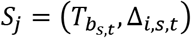. Therefore, each peak *P*_*i*_ with intensity *I*_*i*_ corresponds to a set of subfragments {(*S*_1_, *I*_*i*_), (*S*_2_, *I*_*i*_),…, (*S*_*n*_, *I*_*i*_)}.

### Generating models for the subfragments

To build the prediction models, we collected 34,672 spectra from the GNPS, Mass Bank and in-house databases (Horai et al., 2010; Wang et al., 2016) corresponding to 5,994 unique molecules. For each spectrum in the database, the intensity of the spectrum was normalized so that the sum over all peak intensities equals one. If one molecule has multiple spectra, they will be treated individually. The spectral peaks were converted to fragment substructures as described above. Models were then built for all fragment substructures that occurred in at least five molecules.

To address the imbalance in the number of molecules that do or do not contain a certain fragment substructure in their spectra, individual spectra were assigned weights. For spectra that contained a peak corresponding to the fragment substructure under consideration (positives), the weight is the peak’s intensity. For spectra that did not contain a peak corresponding to the fragment substructure (negatives), equal weights were assigned such that the sum of weights of negatives equals the sum of weights (i.e. normalized peak intensities) of positives.

For each molecule, stereoisomer information was removed, and an 8191-bit chemical fingerprint generated using RDKit Fingerprint. Then, an extra-trees classifier was built using the scikit-learn (Pedregosa et al., 2012) with 100 trees, using the chemical fingerprints of the complete structure as features and the presence of the fragment substructure as class label. A model will be built if a subfragment existed in at least 5 molecules. In total, models were built for 716,447 subfragments. Each of these models predicts the probability *P*(*S*_*i,x*_|*C*) that a peak corresponding to the subfragment *S*_*i,x*_ occurs given the chemical fingerprint *C*.

### Predicting possible peaks for a given molecule

Given a molecule, its chemical fingerprint *C* was calculated as described above. Furthermore, all possible peaks *P*_*i*_ and their corresponding subfragment *S*_*i,x*_ were determined from the molecular structure. The molecule’s chemical fingerprint was then used to predict a probability for the existence of a peak, using the pre-built models for the subfragment. When several subfragment had associated models for a given peak, the highest probability of these was assigned to that peak.

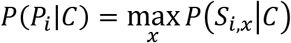

### Spectrum scoring

Given a spectrum with peaks *P*_*i*_ normalized peak intensities *I*_*i*_, the score for a molecule with chemical fingerprint *C* is calculated by summing over the peak probabilities, using the intensities as weights:

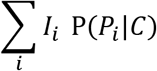

Peaks are determined by searching for molecular formulas within a certain mass accuracy.

### Consensus scoring

For consensus scoring, candidate molecules were scored both with our method as well with CSI:FingerID. A prediction was only accepted if both methods have the same top prediction.

### Performance evaluation

For the CASMI 2016 dataset, the results of CFM-ID and the positive spectra for CSI:FingerID were got from the author’s submission, the results of negative spectra for CSI:FingerID were calculated by Sirius 4.0. For the CASMI 2017 dataset, the results of CSI:FingerID were got from the author’s submission, the results of CFM-ID were calculated by SE-CFM trained model. For the EMBL-MCF dataset, the results of CFM-ID were calculated by SE-CFM trained model, the results of CSI:FingerID were calculated by Sirius 4.0. All the spectra in the test dataset were searched with 20 ppm mass tolerance.

### Data availability and reproducibility

The SF-Matching can be downloaded from http://www.bork.embl.de/Docu/sf_matching. An example of using SF-Matching accessed at https://codeocean.com/capsule/5570439/tree/v1.

## Acknowledgments

The authors thank Joseph Christian Somody, Marco Hennrich, Prasad Phapale and other members of the Bork and Gavin groups as wells as the EMBL Metabolomics Core Facility for helpful discussions, Enric Mila Vilalta for assistance with the benchmark dataset, and Yan Ping Yuan for IT support. We acknowledge funding from EMBL and the MicrobioS grant (ERC-AdG-669830).

## Author Contributions

Methodology, Y.L.; Software, Y.L.; Writing – Original Draft, Y.L, M.K.; Writing – Review & Editing, M.K., A-C.G. and P.B.; Supervision M.K. and P.B.; Funding Acquisition, P.B.;

